# Experimental evolution of Escherichia coli harboring an ancient translation protein

**DOI:** 10.1101/040626

**Authors:** Betül Kaçar, Xueliang Ge, Suparna Sanyal, Eric A. Gaucher

## Abstract

The ability to design synthetic genes and engineer biological systems at the genome scale opens new means by which to characterize phenotypic states and the responses of biological systems to perturbations. One emerging method involves inserting artificial genes into bacterial genomes, and examining how the genome and its new genes adapt to each other. Here we report the development and implementation of a modified approach to this method, in which phylogenetically inferred genes are inserted into a microbial genome, and laboratory evolution is then used to examine the adaptive potential of the resulting hybrid genome. Specifically, we engineered an approximately 700-million-year old inferred ancestral variant of *tufB*, an essential gene encoding Elongation Factor Tu, and inserted it in a modern *Escherichia coli* genome in place of the native *tufB* gene. While the ancient homolog was not lethal to the cell, it did cause a two-fold decrease in organismal fitness, mainly due to reduced protein dosage. We subsequently evolved replicate hybrid bacterial populations for 2,000 generations in the laboratory, and examined the adaptive response via fitness assays, whole-genome sequencing, proteomics and biochemical assays. Hybrid lineages exhibit a general adaptive strategy in which the fitness cost of the ancient gene was ameliorated in part by up-regulation of protein production. We expect that this ancient-modern recombinant method may pave the way for the synthesis of organisms that exhibit ancient phenotypes, and that laboratory evolution of these organisms may prove useful in elucidating insights into historical adaptive processes.

## Background

Understanding historical evolutionary pathways is crucial to understanding how life became the way it is today across millions of years of environmental and ecosystem change (Gould 1989). One of the most difficult aspects of characterizing these historical pathways is the limited amount of knowledge available about how ancient organisms behaved and changed through time. Fossils provide useful morphological and anatomical details, but only traces of information about sub-organismal level processes and states can be inferred from fossilized specimens alone (Pagel 1999). Ancestral sequence reconstruction permits phylogenetics-based sequence inferences of ancestral genes at interior nodes of a tree using likelihood or Bayesian statistics, and offer an opportunity to determine the selectively advantageous amino acid replacements responsible for changes in protein behavior associated with adaptive events for particular molecular systems (Benner 1995; Chang et al. 2002a; Huelsenbeck and Bollback 2001; Liberles 2007; Pauling and Zuckerkandl 1963; Thornton 2004; Ugalde et al. 2004). Going backwards in time at the protein level and studying the biochemical properties of reconstructed ancient proteins in the laboratory may also improve our ability to engineer proteins for specific tasks (Benner 2007; Chang et al. 2002b; Merkl and Sterner 2016; Ogawa and Shirai 2014; Pal et al. 2014). However, mathematical sequence reconstructions and *in vitro* characterization alone may not necessarily provide the salient details of why the protein evolved along a particular evolutionary pathway (Bar-Rogovsky et al. 2015; Copley 2012; Dean and Thornton 2007; Kacar 2016; Zhu et al. 2005), nor does it automatically indicate how the protein’s evolution might be mapped onto the phenotypic evolution of the whole organism, making *a priori* predictions that connect inferred genotype to ancestral phenotype a challenge. Incorporating a functional perspective into the study of ancient proteins may shed light onto the biochemical mechanisms of enzyme evolution, and may also provide clues to the development of ancient adaptive pathways (Dean and Thornton 2007; Harms and Thornton 2013; Kacar and Gaucher 2012; Kacar and Gaucher 2013; Lunzer et al. 2005; Zhu et al. 2005).

Advances in synthetic biology, bioengineering and genomics now allow us to detect changes in genotype and to connect these changes with various perturbations triggered at multiple levels within the cellular environment such as engineering protein-protein interaction networks (Isalan et al. 2008; Wang et al. 2009; Wright et al. 2013), metabolic pathways (Gallagher et al. 2014; Nyerges et al. 2016; Xu et al. 2012), and constructing genetic circuits (Esvelt and Wang 2013; Nandagopal and Elowitz 2011). Further, current methods of evolutionary genome engineering rely upon modifying the genetic content of microbes to execute desired tasks (Andersson et al. 2015; Cambray et al. 2011; Dalchau et al. 2012; Feher et al. 2012; Sandoval et al. 2012; Storz et al. 2015), reducing the genome size to study efficacy of certain proteins and protein families (Annaluru et al. 2015; Deutschbauer et al. 2014; Kolisnychenko et al. 2002; Lee et al. 2005) and heterologously replacing genomic components with homologs obtained from other organisms (Acevedo-Rocha et al. 2013; Agashe et al. 2013; Andersson and Hughes 2009; Pena et al. 2010; Urbanczyk et al. 2012) as well as engineering endogenous promoters with variants obtained across taxa (Nevoigt et al. 2006; Peisajovich et al. 2010). Observing the adaptive pathways of a modified organism can reveal a variety of biochemical pathways adjusting regulatory networks (Dragosits and Mattanovich 2013; Kacar and Gaucher 2012; Michener et al. 2014; Quan et al. 2012), yielding invaluable information on the evolution and prediction of protein function (Counago et al. 2006; Harms and Thornton 2014; Johnsen and Levin 2010; Lind et al. 2010).

An evolutionary bioengineering approach has been proposed to characterize the adaptation of an ancient protein to a modern genome on time scales of laboratory observation; an adaptive laboratory evolution method we previously coined as paleo-experimental evolution (Kacar and Gaucher 2012) (**Figure 1**). It remains to be seen whether it is possible to elucidate and discern ancient adaptive steps from adjustments taken by a modern cell to a maladapted gene. We can begin with a top-level query: if integrated into a modern genome, and given a chance to interact with descendant protein partners in the cell, would the observed mutations alter the ancestral component’s protein structure and function in a manner comparable to the historical pathway inferred for the organism, or will it develop an alternative adaptive pathway? In order to address this, the paleo-experimental evolution system requires an organism with a short generation time and a protein under strong selective constraints in the modern host but whose ancestral genotype and phenotype, if genomically integrated, would cause the modern host to be less fit than a modern population hosting the modern form of the protein. The modern organism hosting the ancient protein would need to be viable, but maladapted. *Escherichia coli* (*E. coli*) and an essential protein family of the bacterial translation machinery, Elongation Factor-Tu (EF-Tu), are ideal for this type of experiment. *E. coli* is an organism that grows quickly in the laboratory, the genetics of the organism are well-known, easy to manipulate, utilizes a range of energy sources and can be stored frozen and later can be re-animated to test ancestral versus evolved populations (Blount 2015). Elongation Factor Tu (bacteria)/elongation factor 1A (archaea and eukaryota) is a GTPase family member involved in the protein translation system (Kavaliauskas et al. 2012). EF-Tu forms a complex with GTP that in turn favors the binding of an aminoacyl-tRNA complex (Agirrezabala and Frank 2009). This ternary complex binds to mRNA-programmed ribosomes thereby delivering aminoacyl-tRNA to the ribosomal A site (Czworkowski and Moore 1996). The biochemistry of EF-Tu has been studied for over three decades giving rise to a clear understanding of the functional aspects of the protein (Negrutskii and El'skaya 1998). Previously, *in vitro* peptide synthesis assays demonstrated that ancestral EF-Tus can participate in a translation system in which all other components necessary for translation besides EF-Tu are provided from modern *E. coli* (Zhou et al. 2012) indicating the possibility that co-evolution between EF-Tu and aa-tRNAs/ribosome/nucleotide-exchange-factors in *E. coli* since the divergence of the ancestral and modern EF-Tu forms has not prevented the ancestral EF-Tu from interacting with the modern *E. coli* translation components. We hypothesized that the diminished capacity of the ancestral EF-Tu is sufficient enough to provide a strong selective constraint for the bacteria containing the ancient gene to acquire beneficial mutations.

**Figure I.**
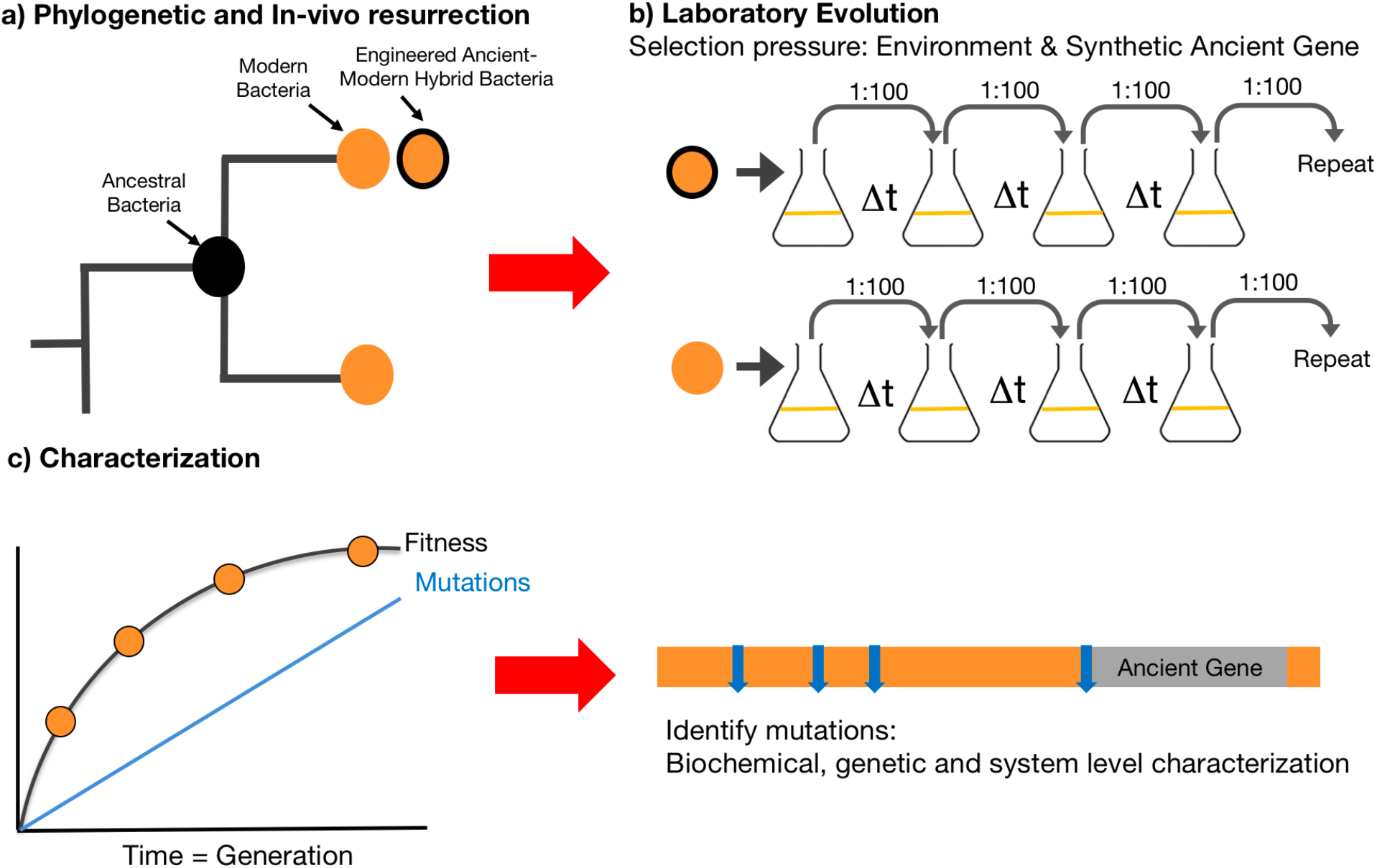
Overall experimental scheme A) Phylogenetic reconstruction allows inferences of the ancestral protein sequence. Replacement of the endogenous EF-Tu gene (orange) with the reconstructed ancient EF-Tu (black) allele is indicated by the black-orange hybrid. B) The hybrid population was subjected to adaptive laboratory evolution through daily propagation and transfer of cultures in minimal media C) Relative fitness of evolved populations were assessed via competitive fitness assays as described by Elena 2005, followed by the identification of mutations through whole-genome sequencing and the characterization of the mutants via in-vivo and in-vitro assays.

Our method builds upon heterologous gene replacement by engineering a synthetic ancient EF-Tu into the genome of *E. coli* bacteria. The engineered EF-Tu represents that of an ancestral γ-proteobacteria that is inferred to be approximately 700 million years old and has 21 (out of 392) amino acid differences with the modern EF-Tu protein (Gaucher et al. 2008). Our experimental system exploits the unique scenario in which *E. coli* bacteria have a paralogous copy of the EF-Tu gene *tufA*, in the form of *tufB*, that frequently recombines with the original copy (Abdulkarim and Hughes 1996). Each of the EF-Tu genes has its own specific expression machinery, and EF-Tu produced through *tufB* accounts for one-third of the cellular EF-Tu as that produced by the *tufA* gene in bacteria (Van Delft et al. 1987; van der Meide et al. 1983; Zengel and Lindahl 1982). Through recombination mediate-engineering (recombineering), the *tufA* gene was deleted from the bacterial genome and the *tufB* copy of a laboratory strain of *E. coli* was replaced with an ancient EF-Tu variant under the control of the endogenous *tufB* promoter, followed by adaptive laboratory evolution of the ancient-modern hybrid populations in replicate lineages through daily propagation of bacterial cultures (**Figure 1**) (Bell 2016; Dragosits and Mattanovich 2013; Elena and Lenski 2003). Evolved populations were sampled for whole genome sequencing, followed by identification of the total number of genomic changes in each population relative to the founding strain and assaying the change in adaptive response through fitness assays. We further investigated whether *in vivo* analyses into the functionality of ancestral components can be used to discern effects arising from the substituted gene when screened from adaptive responses taken by the host cell to the sub-adapted genetic component. Altogether this work provides the first demonstration of an artificial ancient essential gene variant inside a bacterial genome and provides insights into the principles of using experimental evolution for exploring adaptation of artificial genes in modern organisms.

## RESULTS

### Replacement of modern EF-Tu with ancient EF-Tu is detrimental to *E. coli* fitness

Complete replacement of endogenous EF-Tu protein requires disruption of both native *tufA* and *tufB* genes, and insertion of the inferred ancient gene (**Supplementary Figure 1**) (Schnell et al. 2003). We first disrupted the native *tufA* gene. This intermediate *tufA*^−^ *tufB*^+^ construct displays a fitness of 0.89 (*P* < 0.001) relative to the parent clone. Subsequent replacement of *tufB* with the reconstructed ancient *tuf* gene produced a further fitness decline, to 0.77 (*P* < 0.001) relative to the parent clone (**Figure 2**). This dramatic fitness detriment of complete EF-Tu replacement suggests that the ancient gene is compatible with the modern *E. coli* genome, though unfit. Co-evolution between EF-Tu and aa-tRNAs/ribosome/nucleotide-exchange-factors in *E. coli* since the ancestral state for which the ancient *tuf* gene was inferred has thus not prevented the inferred ancestral EF-Tu from interacting with the modern *E. coli* translation system in a viable manner.

**Figure II.**
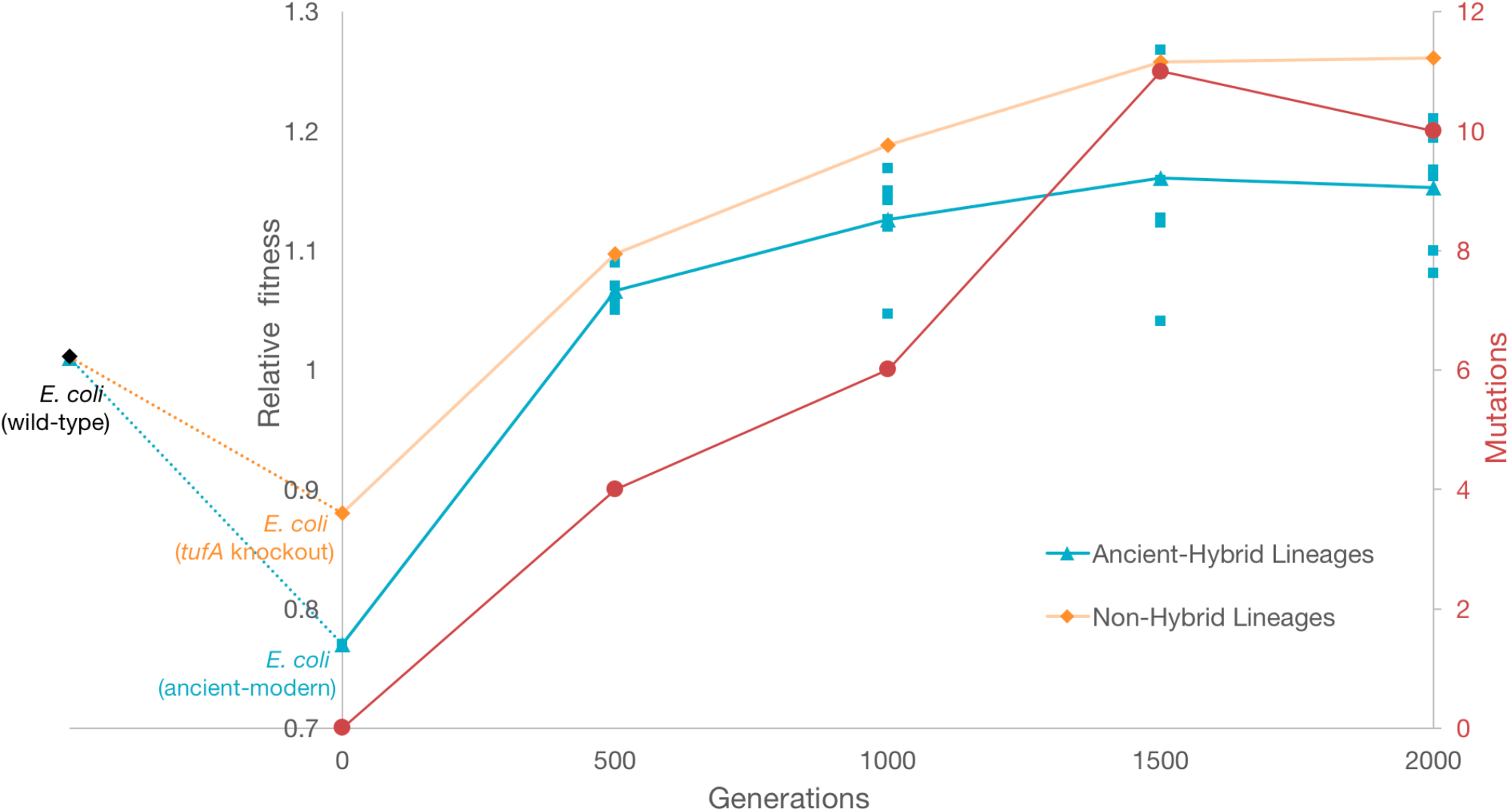
Fitness values of E. coli populations relative to the ancestral strain during adaptive evolution Replacement of the endogenous EF-Tu gene with the reconstructed ancient EF-Tu allele significantly reduces the fitness of the ancient-modern hybrid relative to the original strain (blue dashed line). Hybrid population mean fitness rapidly improved during experimental evolution in minimal glucose medium (blue line). E. coli ΔtufA represents the bacteria that contain a single tufB gene Fitness change in E. coli ΔtufA is shown in orange. Error bars show 95 % confidence interval among six replicate populations for each system. Red line represents the average total number of genomic changes relative to the ancestor in each sampled hybrid lineage. Mutations are total number of genomic changes relative to the ancestor in each sampled lineage that reach to over 20% fixation.

### Experimental evolution allows bacteria to restore fitness

To examine the co-adaptation between *E. coli* and the ancient EF-Tu, we conducted evolution experiments with both the ancient-modern hybrid and the *tufA*^-^ *tufB*^+^ construct. Six replicate populations were generated for each of the two modified genomes by selecting identical clones that were verified to be free of any plasmid vectors that could mediate genetic exchange. The twelve populations were then evolved for 2000 generations under a daily 100-fold serial transfer regime in DM25 minimal glucose medium, at 37°C and 150 rpm. Under these conditions, each population grew log_2_ (100) = 6.64 generations per day before reaching stationary phase. The daily maximum population size for each population is approximately 2.5 × 10^8^ cells. Fitness assays were conducted every 500 generations, in which evolved populations were competed against the ancestral clone. The ancient-modern hybrid populations displayed a mean fitness of 1.06 (*P* <0.043) at generation 500 (**Figure 2**). This increased to 1.12 (P <0.005) at generation 1000, and 1.15 (P<0.009) and 1.16 (P<0.001) at generations 1500 and 2000, respectively. The *tufA*- *tufB*+ populations also exhibited fitness increases. Mean fitness relative to the ancestor is 1.097 (P<0.001) at generation 500, 1.16 (P<0.004) at generation 1000, and 1.15 at both 1500 (P<0.001) and 2000 (P<0.004), respectively (**Figure 2**).

### *Promoter level* mutations upregulate ancient EF-Tu expression and restore bacterial fitness

To identify the genetic bases of the observed fitness increases, whole genomes of clones were periodically isolated and sequenced from all six evolved populations during the evolution experiment. Mutations generally accumulated in similar genes across all experimental construct populations (**Table 1**). However, five out of the six ancient-modern hybrid lineages, but none of the other controls, evolved mutations in the *thrT/tufB* promoter region, with four variant alleles being observed (Lee et al. 1981). The majority of these *thrT/tufB* promoter region mutations accumulated early in the experiment, and rose to high frequency, if not fixation, by 2000 generations across all five populations in which they occurred (**Figure 3**). Such cis-regulatory mutations have been shown to be a common means of adaptation (Hoekstra and Coyne 2007; Jacob and Monod 1961; Lynch and Wagner 2008). Others have observed the accumulation of mutations in orthologous genes engineered into microbes during laboratory evolution experiments. By contrast, we observed no mutations in the ancient or modern EF-Tu gene-coding region in any of the evolved lineages, suggesting that compensatory amino acid replacements may have only occurred at other sites in the genome.

**Figure III.**
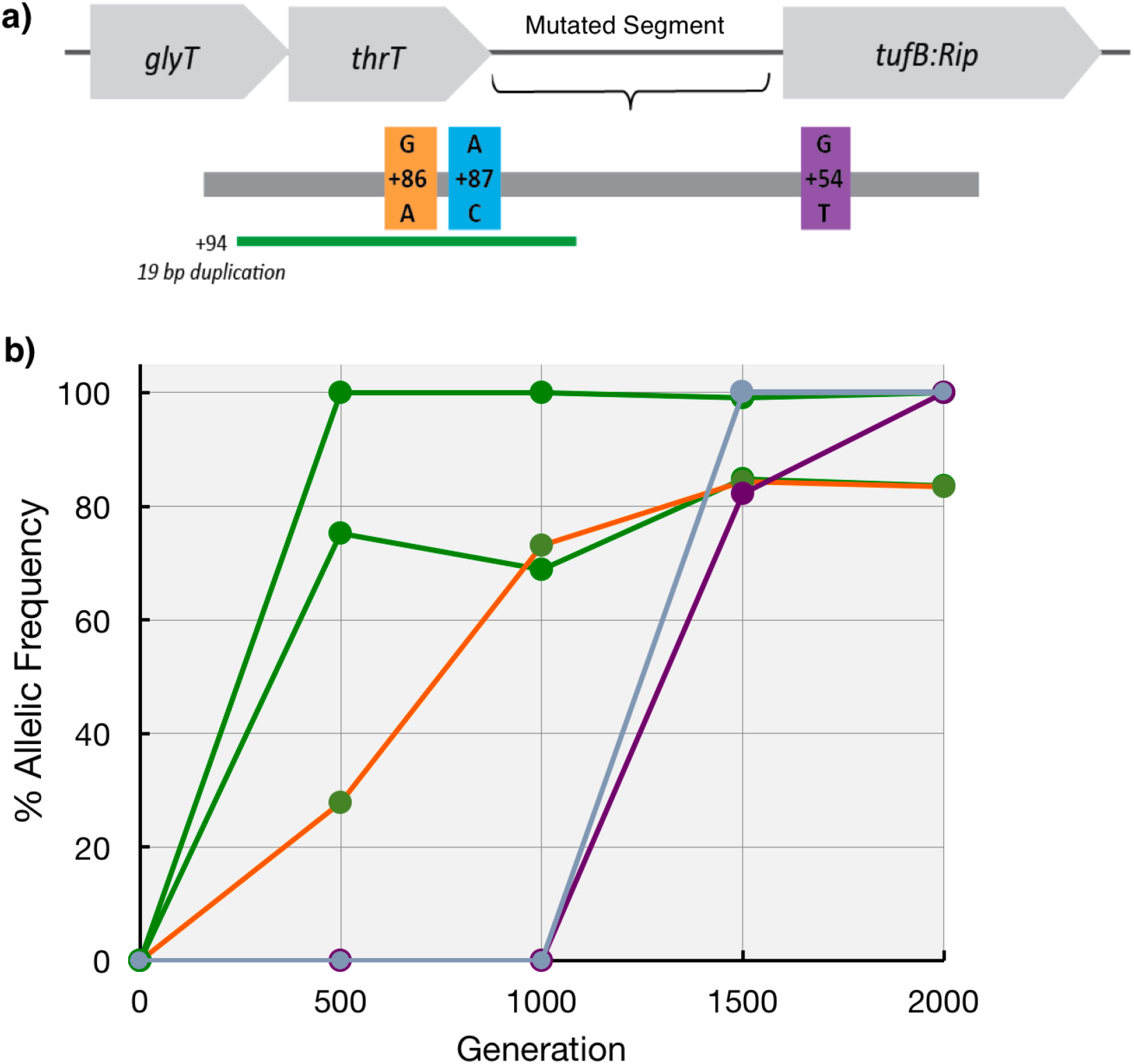
Analysis of the mutations accumulated in the cis-regulatory region thrT/tufB A) The thrT/tufB promoter region in which five of six evolved hybrid populations were found to have accumulated mutations. B) The allelic frequency of the mutations located in ancient EF-Tu gene’s promoter region per generation per population during laboratory evolution.

**Table 1.** Parallel mutations in genes for six, initially identical, independently evolved populations harboring an ancient EF-Tu. Top part represents the genes that accumulated mutations in at least three populations containing the ancient EF-Tu gene and occupied the population by minimum 20% across generation 500 to 2000 are shown for a total of 7 populations evolved in parallel. thrT/tuf represents the intergenic region between ancient EF-Tu gene and thrT gene. Bottom three are the mutated genes that are specific only to the single lineage that did not accumulate a mutation in the thrT/tuf region (referred as Rip 2). Mutations were detected in genes shown with asterisk are the genes in which mutations were detected in at least one population containing the native EF-Tu gene. Prior studies that report mutations in genes reported here include (Barrick et al. 2009, Maddamsetti et al. 2015, Dillon et al. 2016, Conrad 2009, Herron 2013 and Philips et al. 2014)

| Gene | Function | Number of lineages that accumulated a non-synonymous mutation over 20% frequency |  |  |  | Present in non-hybrid lines? | Previously reported for LTEE? |
| --- | --- | --- | --- | --- | --- | --- | --- |
|  |  | Generation |  |  |  |  |  |
|  |  | 500 | 1000 | 1500 | 2000 |  |  |
| mreB | cell wall structural complex MreBCD, actin-like component MreB | 2 | 2 | 3 | 3 | Yes | Yes |
| mrdB | cell wall shape-determining protein | 2 | 2 | 2 | 2 | Yes | Yes |
| thrT/tuf | tRNA-Thr/protein chain elongation factor EF-Tu | 4 | 4 | 5 | 5 | No | No |
| fadA | acetyl-CoA acetyltransferase | 3 | 3 | 3 | 3 | No | Yes |
| ftsZ | cell division protein FtsZ | 1 | 1 | 2 | 5 | No | Yes |
| iclR | DNA-binding transcriptional repressor | - | 1 | 4 | 3 | No | Yes |
| accC | acetyl-CoA carboxylase | - | 1 | 4 | 5 | Yes | Yes |
| pykF | pyruvate kinase | - | 1 | 3 | 5 | Yes | Yes |
| topA | DNA topo isomerase | 1 | 1 | 1 | 1 | Yes | Yes |
| nusA | transcription elongation factor NusA | 1 | 1 | 1 | 1 | No | Yes |
| infB | translation initiation factor IF-2 | 1 | 1 | 1 | - | No | Yes |
| hupA | HU, DNA-binding transcriptional regulator, alpha subunit | 1 | 1 | 1 | 1 | No | Yes |

We performed whole-cell shotgun proteomic analysis on five of the evolved hybrid populations with EF-Tu promoter mutations to examine the impact of these mutations on EF-Tu protein levels. The assayed time points were those for each population at which the mutations had reached over 90% frequency in the population. For comparison, we also assayed unevolved ancient-modern hybrid bacteria, the wild-type parent *E. coli* strain, and an unevolved *tufA*^-^ *tufB*^+^ construct. Deletion of the *tufB* copy and the subsequent insertion of the ancient reconstructed gene into *E. coli* causes EF-Tu protein levels to drop by approximately ~66% relative to that observed in the wild-type. The evolved hybrid populations with *tufB* promoter mutations all show significant increases in EF-Tu levels (**Figure 3, Figure 4a**). We also assessed the effect of these promoter mutations on EF-Tu expression level *in vitro* by examining their effect on a plasmid-borne fluorescent reporter. The mutant promoters increase expression between 1.5 to 20 fold (**Supplementary Figure 2**). Interestingly, promoter mutations that rose to high frequency later during the experiment had lower relative effects on ancient EF-Tu protein expression than those that did so earlier during laboratory evolution. To test whether increased ancient EF-Tu levels would correlate with increased fitness, the unevolved ancient-modern cells, as well as *E. coli* harboring a single wild-type *tufB* gene was transformed with pASK plasmids expressing ancient EF-Tu proteins (Materials and Methods). Overexpression of ancient EF-Tu protein in *E. coli* isogenic strain decreases the doubling time from 35 min to 26 min. Similarly, overexpression of ancient EF-Tu protein in ancient-modern hybrid ancestor decreases the doubling time from 52 min to 29 min (**Figure 4b**). This observation is in agreement with the previous studies demonstrating the correlation between the cellular concentration of EF-Tu and organismal fitness (Brandis et al. 2016; Tubulekas and Hughes 1993). Taken together, these results indicate that each experimental population exhibited parallel patterns of response such as upregulation of EF-Tu, as well as more idiosyncratic means of compensating for altered EF-Tu expression and activity.

**Figure IV.**
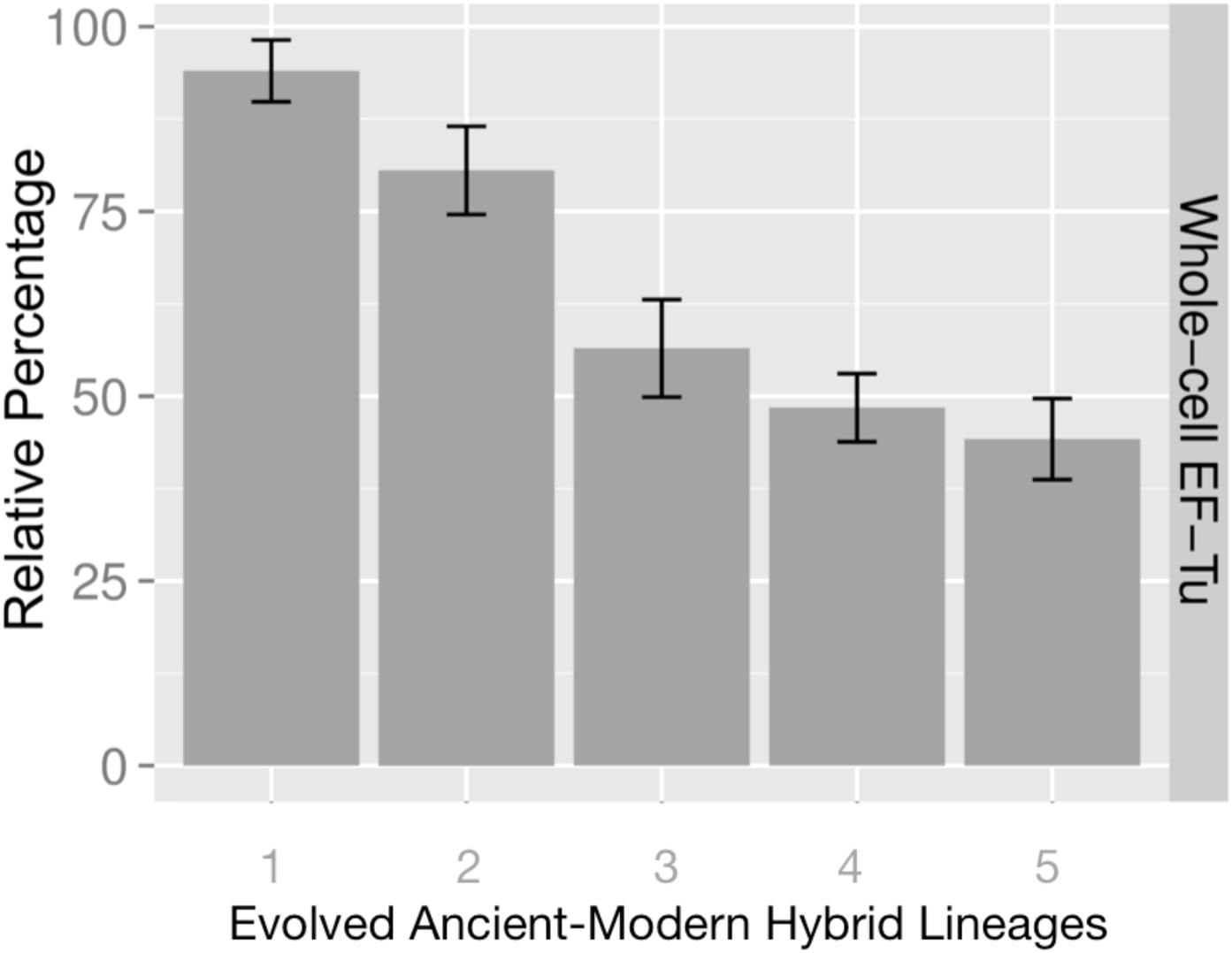
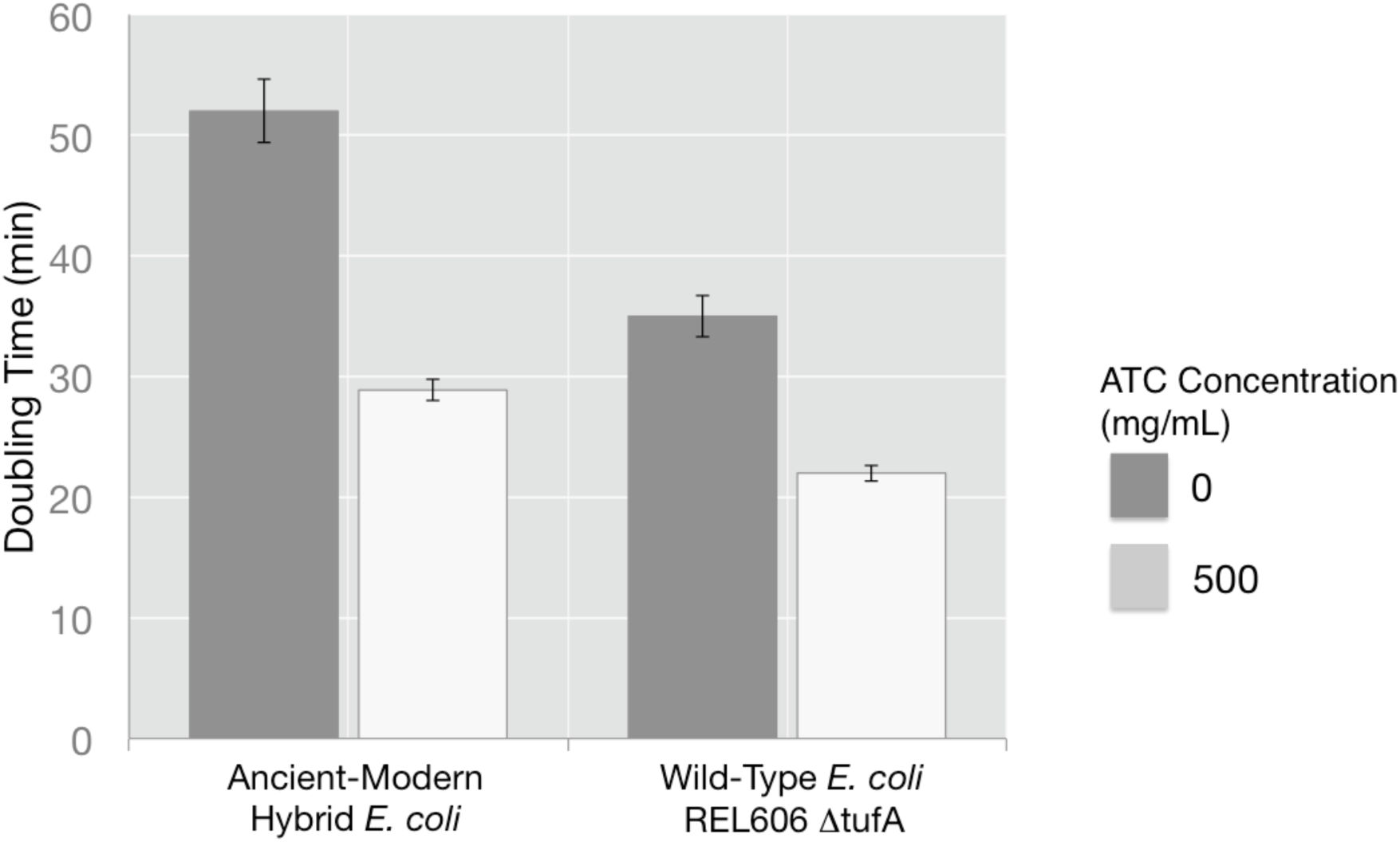
A) Relative abundance of ancient EF-Tu protein among evolved hybrid strains using the peak area quantification from MS proteomics data. Error bars obtained using Anova/t-test. B) Growth rates of an isogenic strain of E. coli REL606 lacking the *tufA* gene, as well as the unevolved ancient-modern hybrid E. coli was evaluated in the presence of with Anhydrotetracycline (ATC) inducer. Strains were induced with 500 mg/mL ATC in rich growth media for 3-4 hours to achieve proper induction. Cells from these fresh induced cultures were inoculated in 96-well plates and grown at 37°C with a starting OD_600_ of −0.06 under respective ATC condition. Doubling times were determined by fitting the exponential growth curves with an exponential function.

### Replacement of the endogenous EF-Tu with the ancient counterpart abolishes previously existing protein level interactions

The single ancient-modern hybrid population in which no *tufB* promoter mutations occurred did accumulate a number of mutations particular to this population, including one in *nusA* gene (**Table 1**). The *nusA* gene is a translation regulator and its protein product is thought to exhibit chaperone activity with direct interaction to ribosomal proteins (Shazand et al. 1993). NusA may affect the efficiency of the translation machinery in a manner similar to EF-Tu and other ribosomal proteins with chaperone activity (Caldas et al. 2000; Caldas et al. 1998). The mutation region is located at the C-terminal domain of the protein, and the complete deletion may be potentially detrimental for the protein’s chaperone activity (**Supplementary Figure 4, 5**). To examine the biochemical effects of the *nusA* mutation, specifically the 27bp deletion (*nusAΔ*9), the mutant protein was cloned in an expression vector and subsequently purified. Changes in interaction of the mutant NusA protein with EF-Tu, ancient or modern was examined by measuring protein-protein binding via isothermal titration calorimetry. While the wild-type EF-Tu bound NusA with a robust binding constant (Kd) of 14.6 ± 5.2 μM, the ancient EF-Tu binds only weakly to the native NusA protein (**Figure 5**). Moreover, dipeptide formation assays detected no NusA-EF-Tu interaction in the ribosome, and the interaction between EF-Tu and NusA had no observable effect on dipeptide formation in the ribosome (**Supplementary Figure 5**).

**Figure V.**
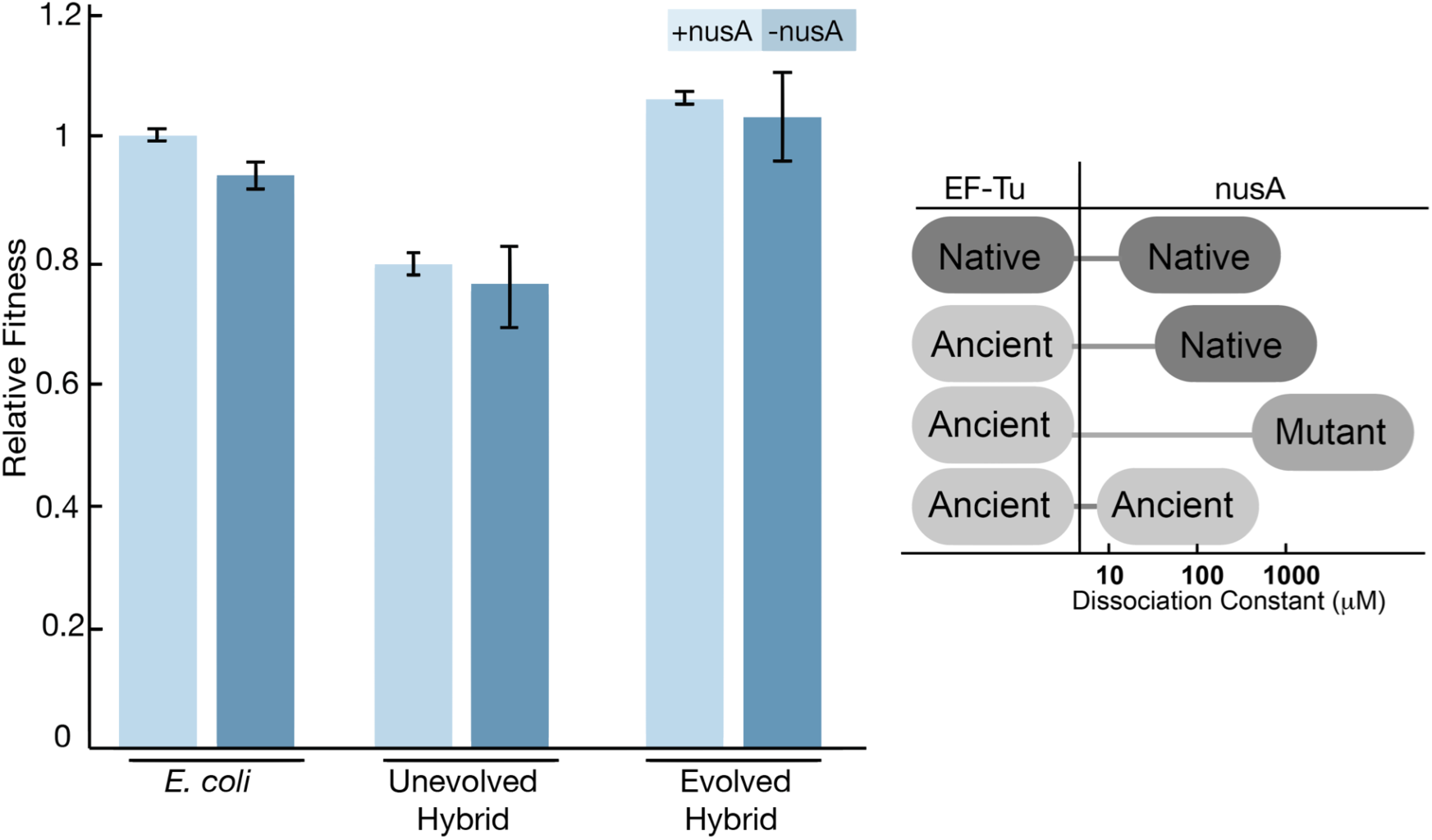
Fitness change after deletion of nusA gene from the ancestral and evolved bacterial genome. (Left) Bacterial constructs with nusA knockouts are constructed and competed against the native E. coli bacteria for fitness measurement. (Right) The interaction between the native EF-Tu, ancient EF-Tu and nusA variants are measured via in vitro Isothermal Calorimetry binding assays.

On the other hand, *nusAΔ*9 and ancient EF-Tu exhibit a Kd of 680 ± 66 μM, suggesting virtually no interaction (**Supplementary Figure 5**). The loss of interaction might be due to the lack of interaction between EF-Tu in the ancestral context in which the reconstructed ancient EF-Tu existed. To test this hypothesis, ancient *nusA* gene representing the γ-proteobacterial ancestor was phylogenetically reconstructed, synthesized, expressed, purified, and examined the ancient NusA protein’s capacity to bind to the ancient EF-Tu protein (**Supplementary Figure 6**). No detectable interaction between the two ancient proteins was observed. Deletion of the mutant *nusA* gene from the evolved and the ancestral allele has no observable fitness effect. However, deletion of *nusA* did cause a statistically significant fitness drop of ~8% in the *tufA*^-^ *tufB*^+^ background (**Figure 5**). The *nusA* mutation is therefore neutral in evolved hybrid background and the mutation is independent of the ancient-EF-Tu mechanism.

### Discussion

By combining a unique set of tools drawn from synthetic biology, evolutionary biology and genomics we experimentally evolved and then analyzed the adaptive properties of a single-celled organism containing a reconstructed ancestral gene inserted in its genome. A majority of the evolved lineages accumulated mutations in the promoter region of the ancestral *tuf* gene and these lead to increased expression of the ancient EF-Tu protein. It is possible that these promoter mutations constitute ‘low hanging fruit’ of compensatory genetic changes, particularly for highly-conserved essential proteins, and that structural mutations in the ancient *tuf* gene might have been observed had evolution been allowed to continue. Understanding the lack of direct accumulation of mutations on the ancient EF-Tu requires a full accounting of the fitness effects of all potentially contributing mutations. Considering the important role of EF-Tu in the translational machinery, mutations accumulating directly on the EF-Tu gene can cause cell lethality and thus may not be readily adaptive (Goldman et al. 2010; Kacar and Gaucher 2013; Pereira-Leal et al. 2006). On the other hand, there likely are beneficial mutations that can occur, but they don't confer the advantage that others do under the same conditions, resulting in tuf mutations always being out-competed early on. Increasing the cellular EF-Tu protein level may be the first emergency response of the organism to cope with a drastic alteration introduced by a maladapted protein central to translation machinery (Bridgham et al. 2009; Gong and Bloom 2014; Kryazhimskiy et al. 2014; Kvitek and Sherlock 2011; Lunzer et al. 2010).

Engineering native genomes with ancient genes has been considered a challenging experimental approach due to the possibility of functional incompatibility of the ancestral genes in modern organisms (Hobbs et al. 2015). Moreover, altering essential genes carries the risk of drastic effects on cellular epistatic networks (Coulomb et al. 2005; Drummond et al. 2005; Zotenko et al. 2008). Even a single mutation in the translation machinery can interfere with protein expression, and thereby drastically impact an organism’s viability (Ito et al. 1998; Lind and Andersson 2013). Indeed, if phenotypically altered by subsequent functional divergence and promiscuity over time (Copley 2003), ancestral sequences could be maladapted to the host cell to the extent that a functional organism is all but precluded. However, this experimental limitation does not apply to reconstructed ancestral genes alone. It has been suggested that as the number of nodes connecting a protein within its protein-protein interaction network increases, the capacity to replace that protein with another homolog decreases despite the presumed functional equivalence between the endogenous gene and the homolog (Jain et al. 1999). While a careful assessment of candidate ancestral protein properties prior to integration is helpful, in most cases, studying gene-triggered genomic perturbations through integration of ancestral genes offers a valuable and complementary alternative to existing methodologies that use orthologous proteins (Pal et al. 2014).

This work was originally conceived as a way to reconstruct the historical pathways by which ancestral genes evolved into modern ones by replacing the modern gene with a reconstructed ancestral one in a modern organism, and then performing experimental evolution with the hybrid. No mutation was detected within the ancestral gene. Instead, all likely adaptive mutations that compensated for the fitness detriment caused by the substitution of the suboptimal ancestral gene occurred at other sites, including within the promoter region of the ancestral gene. This result does not, however, rule out the possibility that useful information on historical gene evolution might be gleaned with this method. The most consistent drivers of historic mutational change may be macroscopic variables (i.e., atmospheric composition, nutrient availability, ecological partitioning, or long-term climate fluctuations) that cannot be readily incorporated into laboratory-scale synthetic evolution experiments. In just one example relevant to this experimental setup, the EF-Tu protein phenotype is tightly coupled to the optimal growth temperature of its host organism (Gromiha et al. 1999), but bacterial clades are not thought to have gone through any significant temperature-dependent evolutionary bottlenecks over the 700 million years of evolution that has occurred between the ages of the modern and ancestral homologous sequences (Blattler and Higgins 2014; Knauth 2004). This is one possible interpretation for the observed lack of mutations on EF-Tu itself. Future applications of this method should focus on linking substituted component behavior with a demonstrable organismal phenotype that can be independently retraced over the inferred age of the component (Kacar et al. 2016).

### Conclusions

Engineering bacterial genomes with phylogenetically reconstructed genes complements the current technique of genome level alterations of gene and gene clusters with currently existing homologs, and provides insights into molecular mechanisms of adaptation by providing access to the historical states of currently existing proteins. However, these methods are also severely constrained by limited existing knowledge of how laboratory evolution setups impact evolutionary trajectories, especially for synthetic biology applications, and the connection between environment and protein phenotype. This knowledge is critical for discerning the change in behavior due to the ancestral state of the protein from the change in systems-level behavior attributable to its intrinsic response to a suboptimal cellular component. The synthetic system described here may enable the development of ancient-modern hybrid model systems that will provide new insights related to the role of evolutionary history and the “tape” of evolution as well as the degree of coupling between protein-level biochemical attributes and macroscale evolutionary trajectories and biogeochemical cycles.

### Acknowledgements

We gratefully acknowledge Zachary Blount, Donald Burke and Seth Childers for their comments on this manuscript. Neerja Hajela for providing the *E. coli* REL606 strain, members of the Richard Lenski laboratory for providing feedback during the earlier phase of this work, Deepak Unni for his assistance during the Breseq analyses, undergraduates Lily Tran (Georgia Tech) and Jennifer Zhang (Georgia Tech) for their assistance in the laboratory, and Brian Hammer (Georgia Tech) for providing the pBBRlux plasmid.

## METHODS

### Bacterial Strains and Culture Conditions

All experiments were done at 37 °C unless stated otherwise. Luria-Bertani (LB) broth was used as the base medium for liquid cultures and agar plates. Experimental evolution and competition assays were carried out in Davis-Mingioli minimal medium (DM) supplemented with 25 mg/L glucose (Davis 1950). Tetrazolium Arabinose (TA) plates were used as the base for competition experiment plating (Reference). When required, LB and DM media were supplemented with kanamycin, chloramphenicol and tetracycline antibiotics. All dilutions were carried out in 0.1% sterile saline. LB and DM cultures were incubated on a rotary shaker 200 rpm and 150 rpm, respectively. The REL606 parent strain was kindly donated by Richard Lenski. Genes coding for ancestral EF-Tu were codon optimized for expression in *E. coli,* chemically synthesized by DNA 2.0, and cloned into a pET15b plasmid as reported previously (Gaucher et al. 2008).

### Recombineering

#### Construction of the ancient-modern hybrid strain

Integration of the ancient EF-Tu gene *(AnEF)* into the chromosome of *E. coli* strain REL606 was carried out via the λ-red homology recombineering approach as described by Datsenko and Wanner (Datsenko and Wanner 2000). First, linear DNA containing homology sequences of upstream and downstream of *tufA* gene was amplified by PCR, via (5’ GTGGTTGCGAAAATCATCGCTAGAATTCCGGGGATCCGTCGACC 3’ and 5’ TGTAATTAGCCCAGAACTTTAGCAACTGTAGGCTGGAGCTGCTTCG 3’), and pKD13 plasmid as template, and then transferred in REL606 cells through electrophoration, together with the temperature sensitive pKD46 plasmid. Recombinants were isolated from LB agar plates containing 50 m g/m l kanamycin at 37 °C, grown in in liquid LB medium containing 50 m g/m L Kanamycin and their genomic DNA was isolated using Promega Wizard Genomic DNA Purification Kit. Confirmation PCR was performed using genomic DNA isolated from colonies as a template, with primers aligning to the chromosome outside of the recombination site (5’ CAGGCCGTAATTGAAGCCCGTGGTAAATAAGCC 3’ and 5’ GAATAATTTATTCGTTCTGACAGTACGAATAAG 3’). Once the successful replacement of *tufA* gene with the kanamycin marker was confirmed via Sanger sequencing, the strain was transformed with linear DNA containing homology sequences of upstream and downstream *tufB* flanked in between the AnEF DNA construct soed to a chloramphenicol marker originally amplified from the A007 loxP-Cm-loxP plasmid (Gene Bridges GmbH) via Gibson Assembly. The transformants were selected on LB plates containing 25 m g/m L Chloramphenicol and 50 m g/m L Kanamycin at 37 °C and the correct insert was screened with primers aligning to the chromosome outside recombination site using Fwd primer 5’ TCCGTGTCTTAGAGGGACAATCGATG 3’and Rev primer 5’ GCAATTAGCTCAGAACTTTTGCTAC 3’. Once confirmed, both the Kanamycin and the Chloramphenicol markers were removed using pCP20 and 706-Cre plasmids (Gene Bridges GmbH), respectively, followed by the confirmation of the deletions by genomic PCR analysis. Plasmids pKD46 and pCP20 were cured by growing the cultures at 42 °C, the final Δ*tufA*, Δt*ufB*:AnEF construct was moved into a fresh ancestral strain via bacteriophage P1 transduction. Freezer stocks of the REL606 *ΔtufA, ΔtufB:AnEF* were prepared by mixing 50% sterile glycerol and overnight liquid cultures originated from a single colony, in 1:2 ratios. All stocks were stored in −80 °C. Isogenic Ara+ variants of the REL606 *ΔtufA, ΔtufB:AnEF* were obtained through gene-gorging protocol (Herring et al. 2003) (plasmid pJEB12 is kindly donated by Jeff Barrick).

#### Deletion of nusA gene from the chromosome

The nusA gene from the chromosome of REL606, ancestral REL606 D *tufA,* D *tufB:AnEF* and evolved REL606 D *tufA,* D tufB:AnEF strain from lineage Rip2 were replaced with a FRT-kan-FRT fragment in the presence of pKD46 helper plasmid as described by Datsenko and Wanner using primers 5’ TCCTGCGTGAAGATATGCTG 3’ and 5’ TCACTTCTTCGCCGATTTCT 3’. PCR amplification of the recombination region and sanger sequencing of this amplified region confirmed the correct replacement and the removal of the selection cassette. The cassette was then removed from the chromosome via pCP20, followed by the curation of pKD46 and pCP20 plasmids at 42°C.

### Growth assays

Saturated overnight cultures were preconditioned by dilution into sterile saline by a 1:100, then again by 1:100 into DM25 medium, followed by an overnight growth. Preconditioned cultures were diluted 1:100 into the assay medium and 100 µL transferred into a 96-well microplate. OD readings were taken at 420 nm every 15 minutes with continuous shaking between readings for 24 hours.

### Experimental Evolution

Experimental evolution was carried out using a serial transfer regime in DM25 medium for 2000 generations (~6.6 generations per day) as described previously (Elena and Lenski 2003). Relative fitness change was measured by competing evolved strains or populations against the ancestral genotype, REL606 or REL607, every 500 generations using a standard competition assay protocol. Relative fitness was defined as the ratio of the Malthusian parameter of one competitor to the other. The Malthusian parameter was calculated as follows: m=(cdx* f^x)/cd0, where cd0 = count of the competitor on day 0, and cdx = count of the competitor on day x, f = growth of the population over time (x-0). In our competitions, f = 100 because our transfers involve 100-fold dilution and subsequent outgrowth.

### Whole genome sequencing

Sequencing libraries of clones of interest were prepared by isolating 3 mg of genomic DNA from bacteria grown in 10 mL LB overnight, fragmented and tagged the isolated DNA with specific Illumina adapters using Nextera DNA sample preparation kit. The product was purified using the Zymo DNA Clean and Concentrator Kit, dual-indexed the libraries with TruSeq Dual Indexed Sequencing primer sets and ensured the products were pure using an Agilent 2100 BioAnalyzer. Sets of compatible barcodes (11plex) were combined into a single lane in an Illumina HiSeq 2500 Rapid Run flow cell (v1) after QC. Sequencing was in a paired end 2 x 100 basepair format (PE100) using TruSeq Rapid SBS reagents. Mutations were identified using the Breseq (0.23) pipeline (Deatherage and Barrick 2014).

#### Fitness measurement of the ancestral strain in the presence of over-expressed EF-Tu

Ancient EF-Tu was cloned into a pASK-IBA43 (IBA Life Sciences) vector inducible under a tetracycline promoter using the following primers Forward 5’ GTTGGAATTCATGTCTAAAGAAAAGTTTGAACGTAC 3’ and Reverse 5’ CGGGATCCTCAAGCGATGATTTTCGCAACCAC 3’, between the Xho and Nde sites leading to plasmid pASK-IBA43-AnEF. Ligation was confirmed using Forward primer 5' GAGTTATTTTACCACTCCCT 3' and Reverse primer 5' CGCAGTAGCGGTAAACG 3'. The plasmid was transferred to REL606 *ΔtufA, ΔtufB:AnEF* cells via electrophoration and selected on LB agar plate with Chloramphenicol. Five representative colonies were picked, preconditioned in LB media containing 250 µM anhydrous tetracycline for 24 hours, followed by a 1:100 dilution into DM media containing glucose. Over-expression of the EF-Tu protein was confirmed through SDS-PAGE gel analysis in comparison to ancestral cells that harbored no plasmid and non-induced plasmid. A REL607 strain was acclimated to the competition environment by separate grown under the same environmental conditions as REL606 *ΔtufA, ΔtufB:AnEF* harboring pASK-IBA43-AnEF. The competitors were then mixed in equal stoichiometric ratios by diluting into fresh DM medium with glucose containing 250 µM anhydrous tetracycline. Samples were plated on tetrazolium arabinose agar plate every 4 hours during the 24 hour competition. The competitions were carried out two times to ensure the precision of fitness estimates.

Whole genome sequencing was completed for 2000 generations for eight lineages harboring ancient EF-Tu, as well as the wild-type strains. To prepare the sequencing library, we isolated 3 mg of genomic DNA from bacteria grown in 10 mL LB overnight, fragmented and tagged the isolated DNA with specific Illimuna adapters using Nextera DNA sample preparation kit. We purified the product using Zymo DNA Clean and Concentrator Kit, dual-indexed the libraries with TruSeq Dual Indexed Sequencing primer sets and ensured the products were pure using a Agilent 2100 BioAnalyzer. We combined the sets of compatible barcodes (11plex) into a single lane on Illimuna HiSeq 2500 Rapid Run flow cell (v1) after QC. Sequencing was in a paired end 2 x 100bp format (PE100) using TruSeq Rapid SBS reagents. The Breseq (0.23) software was used for the generation and the analysis of the mutations (Deatherage and Barrick 2014).

### Fitness measurement of the ancestral strain in the presence of over-expressed EF-Tu

Ancient EF-Tu was cloned into a pASK-IBA43 (IBA Life Sciences) vector inducible under a tetracycline promoter using the primers (Forward) 5’ GTTGGAATTCATGTCTAAAGAAAAGTTTGAACGTAC 3’ and (Reverse) 5’ CGGGATCCTCAAGCGATGATTTTCGCAACCAC 3’, between the *Xho* and *Nde* sites. Ligation was confirmed using Forward primer 5' GAGTTATTTTACCACTCCCT 3' and Reverse primer 5' CGCAGTAGCGGTAAACG 3'. The plasmid was transformed into REL606 *ΔtufA, ΔtufB:AncientEFTu* cells via electroporation, and transformants selected on LB agar plate with chloramphenicol. Five representative colonies were picked, preconditioned in LB media containing 250 µM anhydrous tetracycline for 24 hours, followed by a 1:100 dilution into DM media containing glucose. Over-expression of the EF-Tu protein was confirmed through SDS-PAGE gel analysis in comparison to ancestral cells that harbored no plasmid, or non-induced plasmid. A REL607 strain was acclimated to the competition environment by separate growth under the same environmental conditions as REL606 *ΔtufA, ΔtufB:Rip* harboring pASK-IBA43 with the ancient EF-Tu gene. The competitors were then mixed in 50:50 ratios by volume by diluting each into fresh DM25 supplemented with 250 µM anhydrous tetracycline. Samples were plated on tetrazolium arabinose agar plate every 4 hours during the 24-hour competition. The competitions were carried out twice to increase the precision of fitness estimates.

### Luciferase Assay

#### tufB and pBBRlux plasmid cloning

The wild-type and mutant (evolved) promoter region of *tufB* gene (EF-Tu protein) was cloned into the pBBRlux plasmid as adapted from Lenz et. al (2004) (kindly provided by Prof. Brian Hammer, Georgia Tech). Phusion High-Fidelity DNA polymerase, dNTPs, restriction enzymes (high fidelity), and T4 ligases were all obtained from New England Biolabs. DNA purification materials were purchased from QIAGEN. Promoters were amplified using PCR primers 5’-CAGAATGAAAATCAGGTAGCCGAGTTCCAG-3’ and 5’-TAGTGATTGCAGCGGTCAGCGTTGTTTTAC-3’ and resulted in a 403 bp product from REL606 *E. coli* in the 4155251-4155654 region of the genome. Restriction sites were subsequently added to the ends of the *tufB* promoter with the following primers: Forward 5’-GAT<u>ACTAGTC</u>AGAATGAAAATCAGGTAGCCGAGTTCCAG-3’ and Reverse 5’-TAT<u>GGATCC</u>TAGTGATTGCAGCGGTCAGCGTTGTTTTAC-3’ (underlying restriction sites correspond to *SpeI* and *Bam*HI, respectively). The EF-Tu promoter was cloned upstream of the luciferase operon in the pBBRlux plasmid in order to drive transcription. pBBRlux provides chloramphenicol (CMP) resistance.

#### Scintillation Counts

Four experimental constructs: +86/-29 (G+86A), +54/-61 (G+54T), +87/-28 (A+87C), +94/-21 (19bp duplication, +96); and two control constructs: P (no promoter), Patuf (wild-type, or unevolved ancestor, *tufB* promoter) were transformed into chemically competent *E. coli* (REL606) cells and incubated at 37 °C for 24 hours on chloramphenicol (CMP) agar plates. A single colony was cultured in LB media containing CMP at 37°C for 24 hours. A 100µL aliquot of the overnight culture was diluted one thousand-fold prior to being transferred into a 50 mL Erlenmeyer flask containing 9.9 mL of DM25 media. Cells were grown for ~8.25 hours, or ~5 doublings as monitored by plating (this represents the end of log growth since these cultures reach stationary phase after ~6.6 generations in DM25) and then pelleted. The supernatant was aspirated until 100 µL of media remained, and the pellet was then resuspended in the remaining 100 µL. Scintillation counting was used to quantify the amount of light signal generated by the luciferase pathway. For all six constructs, three readings per sample were averaged for each of the two replicates assayed.

### Bacterial Enumeration

For each construct, a 10µL aliquot was serially diluted 50 thousand-fold and 50 µL was plated on agar petri dishes containing CMP. Extrapolation was utilized to determine the total amount of cells in each scintillation assay. Three plates per flask were averaged.

#### Luciferase Assay Statistical Analysis

The luciferase expression per cell was normalized by:

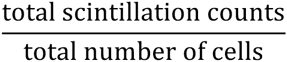

Luciferase expression for each construct was subtracted by the amount of luciferase signal from P to eliminate any leaky expression from the pBBRlux vector without promoter and presented as fold-change relative to the amount of luciferase signal from Patuf. A one-way ANOVA with α = 0.05 and a post-hoc Tukey HSD Test were performed against Patuf to determine significant differences.

### Protein biochemistry

#### Cloning, expression and purification of modern EF-Tu and ancient EF-Tu proteins

Both of the EF-Tu genes were ligated into pET15b plasmid between BamH1/EcoR1 sites, containing an N-term His-Tag with Ampicilin resistance. For expression, the plasmids were transferred in a BL21(DE3) strain, the cells were grown in LB media until OD_600_ reading reached 0.6-0.8 and then induced with 1 mM Imidazole for 4 hours. The cells were lysed using Bugbuster protein extraction reagent (EMD Millipore) containing benzonase. For purification of the His-tagged protein from the supernatant, the cleared lysate was transferred into nitrilotriacetic acid (Ni-NTA) resin gravity-flow columns (Qiagen, Hilden, Germany) at 4°C that was pre-equilibriated with lysis buffer (50 mM NaH_2_PO_4_, 300 mM NaCl, 10 mM imidazole, pH 8). The Ni-NTA gravity-flow column was washed by two times wash with lysis buffer containing 20 mM imidazole. His-tagged protein was eluted using elution buffer (50 mM NaH_2_PO_4_, 300 mM NaCl, 200 mM imidazole, pH 8).

#### Cloning, expression and purification of NusA proteins

Both the wild type *nusA* and the evolved *nusA* genes were amplified from their host bacterial genome using Forward primer 5'-GTGAAGGTGTCGACGCTGCGTGCGCT-3' and Reverse primer 5'-AGCGCACGCAGCGTCGACACCTTCAC-3'. The amplified DNA was purified through gel extraction, removed from salt and then cloned into a pET15b vector (Novagen) using 5’ GGCGACATATGAACAAAGAAATTTTGGC 3’ and 5’ GGAGCTCGAGTTACGCTTCGTCACCGA 3’ primers in between BamH1 and XhoI sites. The plasmids were transferred into a BL21(DE3) strain for expression and induced by IPTG. Cells were broken by French Press in Buffer A (20 mM Tris-HCl at pH 7.5, 50 mM MgCl_2_, 200 mM NaCl, 5% glycerol). 25 μM GDP was added in Buffer A for EF-Tu purification. After centrifugation for 30 min at 16,000 rpm (F21-8x50 rotor, Thermo), the supernatant was applied to Ni-NTA column and elute with gradient Buffer B (Buffer A supplied with 500 mM imidazole). To prepare EF-Tu and NusA for ITC experiment, the proteins were dialyzed in Buffer C (20 mM Tris-HCl at pH 7.5, 50 mM MgCl_2_, 100 mM KCl) for 16 hours at 4 °C.

#### ITC Analysis

The ITC data was measured on a Microcal ITC200 System (GE Healthcare). The syringe was loaded with 42 μL of 0.6-1 mM NusA and the sample cell was filled with 10uM EF-Tu. NusA was titrated (2.5 μL for each) into EF-Tu with 120 s intervals and the first injection was 0.25 μL. The stirring speed was set at 1000 rpm. Blank experiment was measured by titrating NusA into Buffer C (20 mM Tris-HCl at pH 7.5, 50 mM MgCl_2_, 100 mM KCl).

### LC-MS/MS Analysis

#### Sample preparation

Whole cell lysate was generated from each ancestral and evolved strains using Bug Buster reagent (EMD Millipore) and following manufacturer’s instructions. Total protein was quantified via BCA assay using Pierce BCA protein assay kit (Thermo Fisher Scientific). 30 mg of whole cell lysate were submitted to the Proteomics and Metabolomics Facility at Colorado State University. Samples were processed for in-solution trypsin digestion as previously described (Schauer et al. 2013). Briefly, protein was precipitated out of solution in the presence of 4 volumes of 100% −20°C acetone and then resolubilized in 8 M urea, 0.2% ProteaseMAXtm surfactant trypsin enhancer (Promega, Madison, WI). Samples were reduced and alkylated with 5mM dithiothreitol and 5 mM iodoacetamide. Trypsin (MS Grade, Thermo Pierce, San Jose, CA) was added at an enzyme to substrate ratio of 1:50 and incubated at 37 °C for 3-hours. Trypsin was deactivated with the addition of 5% trifluoroacetic acid and desalted using C18 OMIX tips (Agilent Technologies, Santa Clara, CA) using manufacturer’s instructions. Peptide eluate was dried in a vacuum evaporator and resuspended in 3% acetonitrile/0.1% formic acid at a concentration of approximately 1 m g/m l. Relative Quantitation of EF-Tu proteins were carried out using spectral counting approach. Approximately 2 m g of tryptic digest for each sample was injected using an EASY nanoLC-II system (Thermo Scientific, San Jose, CA). Peptides were purified and concentrated using an on-line enrichment column (EASY-Column, 100 m m ID x 2cm ReproSil-Pur C18). Subsequent chromatographic separation was performed on a reverse phase nanospray column EASY-Column, 3 m m, 75 m m ID x 100mm ReproSil-Pur C18) using a 180 minute linear gradient from 10%-55% buffer B (100% ACN, 0.1% formic acid) at a flow rate of 400 nanoliters/min. Peptides were eluted directly into the mass spectrometer (Thermo Scientific Orbitrap Velos). The instrument was operated in Orbitrap-LTQ mode where precursor measurements were acquired in the Orbitrap (60,000 resolution) and MS/MS spectra (top 20) were acquired in the LTQ ion trap with normalized collision energy of 35%. Mass spectra were collected over a m/z range of 400-2000 Da using a dynamic exclusion limit of 2 MS/MS spectra of a given peptide mass for 30 s (exclusion duration of 90 s). Compound lists of the resulting spectra were generated using Xcalibur 2.2 software (Thermo Scientific) with a S/N threshold of 1.5 and 1 scan/group.

#### Data Analysis – Spectral Counting

*Database searching* Tandem mass spectra were extracted, charge state deconvoluted and deisotoped by ProteoWizard version 3.0. All MS/MS samples were analyzed using Mascot (Matrix Science, London, UK; version 2.3.02). Mascot was set up to search the Uniprot_e_coli_custom_reverse database (Updated August 2014, 8750 entries) (Elias and Gygi 2010) assuming the digestion enzyme trypsin, allowing up to 3 missed cleavages. Mascot was searched with a fragment ion mass tolerance of 0.80 Da and a parent ion tolerance of 20 PPM. Oxidation of methionine M (+15.99) and carbamidomethyl of cysteine C (+57) were specified in Mascot as variable modifications.

#### Criteria for protein identification

Scaffold (version Scaffold_4.3.4, Proteome Software Inc., Portland, OR) was used to validate MS/MS based peptide and protein identifications. Peptide identifications were accepted if they could be established at greater than 69.0% probability to achieve an FDR less than 0.1% by the Scaffold Local FDR algorithm. Protein identifications were accepted if they could be established at greater than 99.0% probability to achieve an FDR less than 1.0% and contained at least 2 identified peptides (Kall et al. 2008; Keller et al. 2002). Protein probabilities were assigned by the Protein Prophet algorithm (Nesvizhskii et al. 2003). Proteins that contained similar peptides and could not be differentiated based on MS/MS analysis alone were grouped to satisfy the principles of parsimony.

#### Quantitative analysis

Binary comparisons were created in separate Scaffold files comparing wild type *E. coli* REL606 and unevolved ancestor harboring the ancient protein and the evolved lineages tested (biological replicates n=3) to Strain/Treatment Group (each n=3). Biological samples were organized into Categories based on strain type. Each Category had 3 biological replicates. Normalization of spectral counts was not applied based on these criteria: An equal amount of sample from each replicate was loaded into the mass spectrometer and there was no deviation in processing and the number of spectra between samples is closely similar (% CV < 5% between biological replicates). Spectral counting uses the sum of the MS/MS spectra assigned to each protein as a measure of abundance (Paoletti and Washburn 2006). A T-Test was performed on Total Spectral Counts for each MS sample using the embedded algorithm in Scaffold v 4.3.4. Proteins with P-values less than 0.05 are excluded in calculation of fold changes compared to *E. coli* REL606.

## Supplementary Material Captions

**Supplementary Figure I**

Recombineering outline for the allelic replacement of the tufA with the ancient EF-Tu gene

**Supplementary Figure II**

A) Effect of mutations on the expression of reporter gene for the EF-Tu promoter region measured using luciferase reporter assay. B) Relative whole cell concentration of EF-Tu (percent) based on MALDI TOF MS detection

**Supplemental Figure III**

The replacement of the modern EF-Tu with the ancient EF-Tu caused E. coli to be maladapted. As the ancient-modern hybrid populations evolve, the doubling time reduces from ~70 minutes to ~45 minutes. Each red line represents the mean doubling time calculated via three representative clones for each lineage (n=7) in minimal glucose media (DM25).

**Supplemental Figure IV**

ITC profiles for the titration of NusA to EF-Tu. NusA protein that was purified from plasmids pLT20 (wild-type nusA) (a and c), pLT21 (mutant nusA with 27bp deletion in C-terminal, nusA ΔCTD27) (b and d) was injected to 10 μM wild type (a and b) and ancient EF-Tu (c and d), respectively. The upper panel of each figure represents the raw plots of enthalpy for each injection (μcal/s) against time (min). The corresponding bottom panels show integrated heats (closed squares) in each injection against mole ratio. The data points were fitted to a one-site model, suggesting that native EF-Tu interacts with nusA with a Kd of 14.6 ± 5.2 μM.

**Supplemental Figure V**

A) SDS PAGE analysis of the ribosomal elongation complex shows that NusA cannot bind to 70S ribosome. Initiation complex (containing 70S ribosome, initiation factors Kd (μM) Wild type NusA NusA Δ9 Wild type EF-Tu 14.6 ± 5.2 153.6 ± 5.4 Ancient EF-Tu 30.1 ± 2.6 680 ± 66 (IF1, IF2 and IF3), XR7-ML mRNA and fMet-tRNAfMet) was mixed with elongation mix (containing EF-Tu GTPase mutant H84A, EF-Ts, Leu tRNA synthetase, Leucine) in the absence and presence of NusA. After incubating the reactions for 10s the mixes were loaded directly on 37% sucrose cushion (100 ml). The samples were centrifuged at 80,000 rpm for 2 hours at 4°C. Ribosomal pellets were loaded in the SDS-PAGE. The gel was stained with Coomassie blue and only the upper part of the gel showing bands corresponding to 30S protein S1, EF-Tu H84A and NusA is displayed. EF-Tu H84A was chosen to show EF-Tu retention in the ribosome since this mutant EF-Tu is defective in release.

The lanes contained following samples:

1. 70S initiation complex (IC), 2. Elongation mix (EM), 3. EF-Tu H84A protein,

4. 70S IC + EM + NusA, 5. NusA protein alone,

6. 70S IC + EM + NusAD27CTD, 7. NusAΔ9 alone,

8. 70S IC + EM, 9. 70S ribosome

The absence of NusA bands in lanes 4 and 6 suggest the lack of binding of NusA to the 70S ribosome.

**Supplementary Figure VI**

NusA protein structure highlighting the two alleles observed during the adaptive laboratory evolution. Nus A protein is made up of a dual repeated acidic domain of 70 residues. These modules are referred to as acidic repeats (AR) 1 and 2. Mutations are located in the C-Terminal, away from the its N-protein binding site.

**Supplementary Figure VII**

Sequence alignment of the reconstructed ancestral and present-day NusA protein.

